# Enhancer/Enhanceosome/Super-enhancer and the promoter-proximate RNA Polymerase II (Pol II) pausing are coupled

**DOI:** 10.64898/2025.12.10.693453

**Authors:** Haolin Liu, Lingbo Li, Qianqian Zhang, Christopher Ott, Gongyi Zhang

## Abstract

Virus infection triggers the expression of IFN-xβ, which requires at least four groups of transcription factors and p300/CBP to form an enhanceosome. The promoter-proximal Pol II pausing regulation in metazoans includes the recruitment of CDK9 by JMJD6 and BRD4, the generation of a unique phosphorylation pattern of CTD of Pol II, the recruitment of JMJD5, and the cleavage of arginine methylated histone tails on the +1 nucleosome. We find that both BRD4 and p300/CBP control the activity of CDK9. It is likely that the BRD4 docking site on nucleosomes is generated by p300/CBP. We conclude that enhancer/enhanceosome/super-enhancer and Pol II pausing regulation are coupled and could be the unique universal mechanism that differentiates animals from unicellular organisms.

---

It was first reported that human IFN-β gene expression after virus infection requires the assembly of an enhanceosome, which includes four groups of transcription factors, the high mobility group protein HMGI, ATF2/c-JUN, NF-κB, and interferon response factors (IRFs), respectively (Thanos and Maniatis 1995). p300/CBP was found to be tightly associated with the enhanceosome and to be essential for the expression of IFN-β (Merika, Williams et al. 1998, Wathelet, Lin et al. 1998). Detailed characterization showed that NF-κB is first recruited less than 2 hrs after virus infection, followed by IRF-1 (2-3 hrs after infection), p300/CBP (2-3 hrs), ATF2 (3-4 hrs) (Agalioti, Lomvardas et al. 2000, Lomvardas and Thanos 2002). Interestingly, RNA Polymerase II (Pol II) is recruited at the promoter region of the IFN-β gene around 4 hrs after virus infection while mRNA of IFN-β was detected 6 hrs after virus infection (Agalioti, Lomvardas et al. 2000, Lomvardas and Thanos 2002). It takes almost two hours for Pol II to be ready for transcription after recruitment, suggesting that the participation of Pol II pausing mechanism in the process, which was identified as a unique transcription regulation of Pol II in metazoans (Gilmour and Lis 1986). Interestingly, after deep characterization of the underlying mechanism of the function of the enhanceosome, it was found that the +1 nucleosome located from transcription start site of IFN-β gene was gone though the authors interpreted that the +1 nucleosome was sliding downstream after virus infection via chromatin remodelers such as SWI/SNF family and proposed that the enhanceosome is required to expose the promoter region for Pol II to bind (Lomvardas and Thanos 2001). However, SWI/SNF (BRG1) was recruited 6 hrs (similar time point of appearance of mRNA of IFN-β gene) after virus infection while Pol II was recruited 2-3 hrs earlier (Agalioti, Lomvardas et al. 2000, Lomvardas and Thanos 2002), which is consistent with the result from the same group showed that Pol II was recruited to transcribe even without the enhanceosome in vitro (Lomvardas and Thanos 2001). Another interesting observation from the report was that p300/CBP remains at the promoter region while both SWI/SNF (BRG1) and GCN5 disappear quickly (Agalioti, Lomvardas et al. 2000). The correspondent PI proposed that chromatin remodelers are drivers to carry histone modification complexes along gene bodies to carry out correspondent modifications, such as SWI/SNF drives GCN5 complex (SAGA) to carry out pan-acetylation on histone tails of H2A and H3 along gene bodies (Zhang 2024). This is consistent with the transient existence of both SWI/SNF and GCN5 at the promoter region of IFN-β gene. In conclusion, it is likely that the enhanceosome participates in the depletion or ejection of the +1 nucleosome at the promoter of the IFN-gene to trigger the release of paused Pol II instead of exposing the promoter region to recruit Pol II during the early stage of virus infection.

Since the discovery of the special transcription regulation in metazoans (Gilmour and Lis 1986), the underlying mechanism or the exact functional role of promoter-proximal Pol II pausing remains a mystery. A long-standing view is that Pol II pausing is initiated by Spt4/5 (Wada, Takagi et al. 1998) and NELF (Yamaguchi, Takagi et al. 1999). However, several groups showed that +1 nucleosomes are the barriers, which lead to the pause of initiated Pol II (Mavrich, Jiang et al. 2008, Schones, Cui et al. 2008, Weber, Ramachandran et al. 2014). We surreptitiously discovered that JMJD5/JMJD7 specifically cleave arginine methylated histone tails to generate tailless nucleosomes, which could be overcome by Pol II due to the compromised interaction between DNA and tailless histone octamer (fragile nucleosome, ejected by Pol II) and the subsequent release of Pol II from the status of pausing (Liu, Wang et al. 2017, Liu, Wang et al. 2018). Further characterization revealed that arginine methylated +1 nucleosome is the main target of JMJD5, which was recruited by a unique phosphorylation pattern of the C-terminal domain (CTD) of Rpb1 of Pol II via the CTD interaction domain (CID) within JMJD5 (Liu, Ramachandran et al. 2020). Interestingly, this unique phosphorylation pattern (Ser2ph-Ser2ph-Ser5ph-CTD of Pol II) was specifically generated by CDK9 in metazoans (Liu, Ramachandran et al. 2020). Moreover, we found that JMJD6 cleaves MePCE to disrupt the 7SK snRNP complex to help BRD4 recruit CDK9 onto Pol II, a similar mechanism used by the Tat protein to hijack the CDK9 complex for the activation of the HIV genome (Lee, Liu et al. 2020). It was well established that CDK9 is essentially required to release paused Pol II in metazoans (Marshall and Price 1995), while the +1 nucleosomes for a large group of genes in metazoans cause the pause or stalking of Pol II (Mavrich, Jiang et al. 2008, Schones, Cui et al. 2008, Weber, Ramachandran et al. 2014). Putting all these data together, a general underlying mechanism of Pol II pausing regulation in metazoans could be derived (Zhang 2024).

From the study of the expression of IFN-β gene, Pol II was paused at the promoter region of IFN-β gene. Enhanceosome is required to remove the +1 nucleosome to trigger the expression of IFN-β gene. Results from our group showed that arginine methylated histone tails on the +1 nucleosome are cleaved by JMJD5/JMJD7, which in turn triggers the release of paused Pol II. Two independent story series suggest that the enhanceosome and the paused Pol II regulation in metazoans could be coupled. To prove this hypothesis, we reason that the activities of p300/CBP could be the critical component to bridge the two stories. It was reported that p300/CBP is essential to generate acetyl-groups on the histone tails of nucleosomes at enhancer/promoter regions to generate docking sites to recruit BRD4 (Jin, Yu et al. 2011, Zhang, Prakash et al. 2012). Furthermore, it was reported that p300/CBP and Pol II pausing are coupled (Narita, Ito et al. 2021). Our data showed that BRD4, with the help of JMJD6, can recruit CDK9 onto Pol II (Lee, Liu et al. 2020). To confirm the role of BRD4 in the recruitment of CDK9, we applied a PROTAC inhibitor of BRD4, dBet6, which specifically degrade BRD4 via binding to the bromodomain of BRD4 to check the change of the activity of CDK9, the level of Ser2ph-Ser2ph-Serp5-CTD of Rbp1 of Pol II, an accurate readout assay for the activity of CDK9 established by us (Lee, Liu et al. 2020, Liu, Ramachandran et al. 2020). To our satisfaction, incubation of dBet6 leads to the drop of Ser2ph-Ser2ph-Serp5-CTD of Rbp1 of Pol II after 6 hrs at 250 nM (**Fig. 1A**). If p300/CBP is required to generate docking sites to recruit BRD4, we expect that inhibition or acute removal of p300/CBP will also lead to failure of the recruitment of BRD4 and subsequent the drop of Ser2ph-Ser2ph-Serp5-CTD of Rpb1 of Pol II. dCBP-1, a potent p300/CBP degrader, was incubated with HEK293T cells, p300/CBP was almost gone with the incubation of dCBP-1 after 6hrs at 10 μM (**Fig. 1B**). Interestingly, the level of Ser2ph-Ser2ph-Serp5-CTD of Rpb1 of Pol II is almost all gone too. The result supports that both BRD4 and CBP/p300 work upstream the function of CDK9.

**Figure 1.**
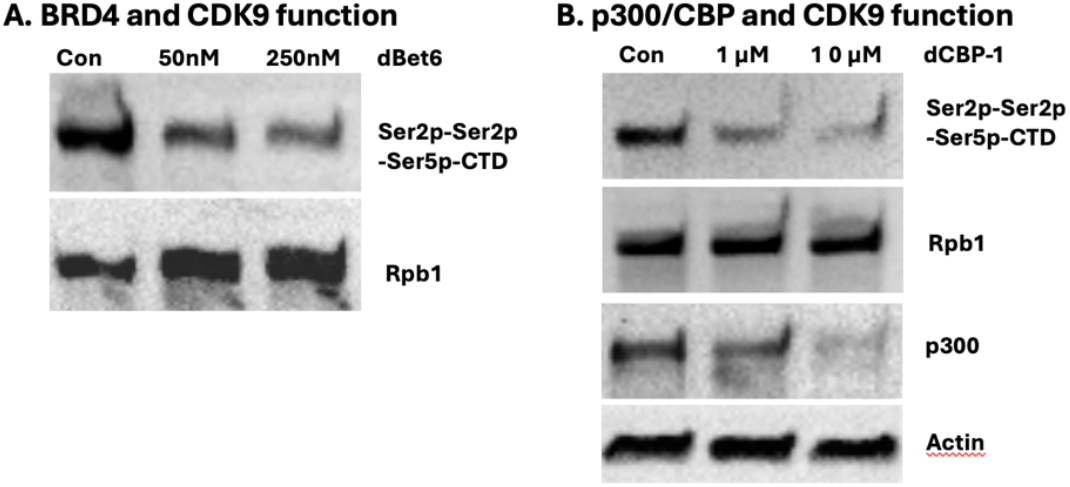
P300/CBP is essential to generate a docking site to recruit BRD4, which works upstream of the CDK9 complex. **A**. Acute removal of BRD4 leads to the drop of the unique phosphorylation pattern of CTD generated by CDK9. **B**. Acute removal of p300/CBP also leads to the disappearance of the unique phosphorylation pattern of CTD.

The enhanceosome was identified three decades ago (Giese, Kingsley et al. 1995, Thanos and Maniatis 1995). The underlying mechanism of this synergic action of a group of transcription activators remains ambiguous. With the elucidation of the underlying mechanism of the promoter-proximal Pol II pausing regulation, we finally found that enhanceosome and the Pol II pause regulation are coupled and bridged by p300/CBP and BRD4, which are characteristic components for a super-enhancer and could be the same content with different manifestations. Based on our new data and others, a final model could be derived (**Fig. 2**). Virus infection triggers the activation of three groups of transcription factors, NF-κB, IRFs, and ATF-2/c-JUN via phosphorylation and exposing activation motifs of the three groups of transcription factors, which are usually buried in condensates formed by intrinsically disordered regions-IDRs (Boija, Klein et al. 2021). Activated transcription factors will migrate into the nucleus and bind to the enhancer region of the IFN-β gene, which was maintained (kept the enhancer exposed without nucleosome) by HMGI. The three groups of transcription factors will recruit p300/CBP via exposed activation motifs to form the enhanceosome or super-enhancer. p300/CBP, in turn, acetylates the histone tails on the −1 nucleosome, which recruits BRD4. BRD4, with the help of JMJD6, recruits the CDK9 complex onto the super-elongation complex (SEC), which could be recruited onto Pol II by ELL2. CDK9 complex then phosphorylates the CTD of Rpb1 to generate the docking sites to recruit JMJD5. JMJD5 cleaves the arginine methylated histone tails on +1 nucleosome to generate a fragile tailless nucleosome. Initiated Pol II will overcome the fragile +1 nucleosome and get into the mode of elongation (**Fig. 2**). Based on the commonality of enhanceosome/super-enhancer and promoter-proximal Pol II pausing regulation in the development of animals but lacking in simple unicellular organisms, it is likely that this coupled regulation mechanism may participate in all genes involved in development, differentiation, and environmental stimulus, and consequently differentiate animals from simple unicellular organisms.

**Figure 2.**
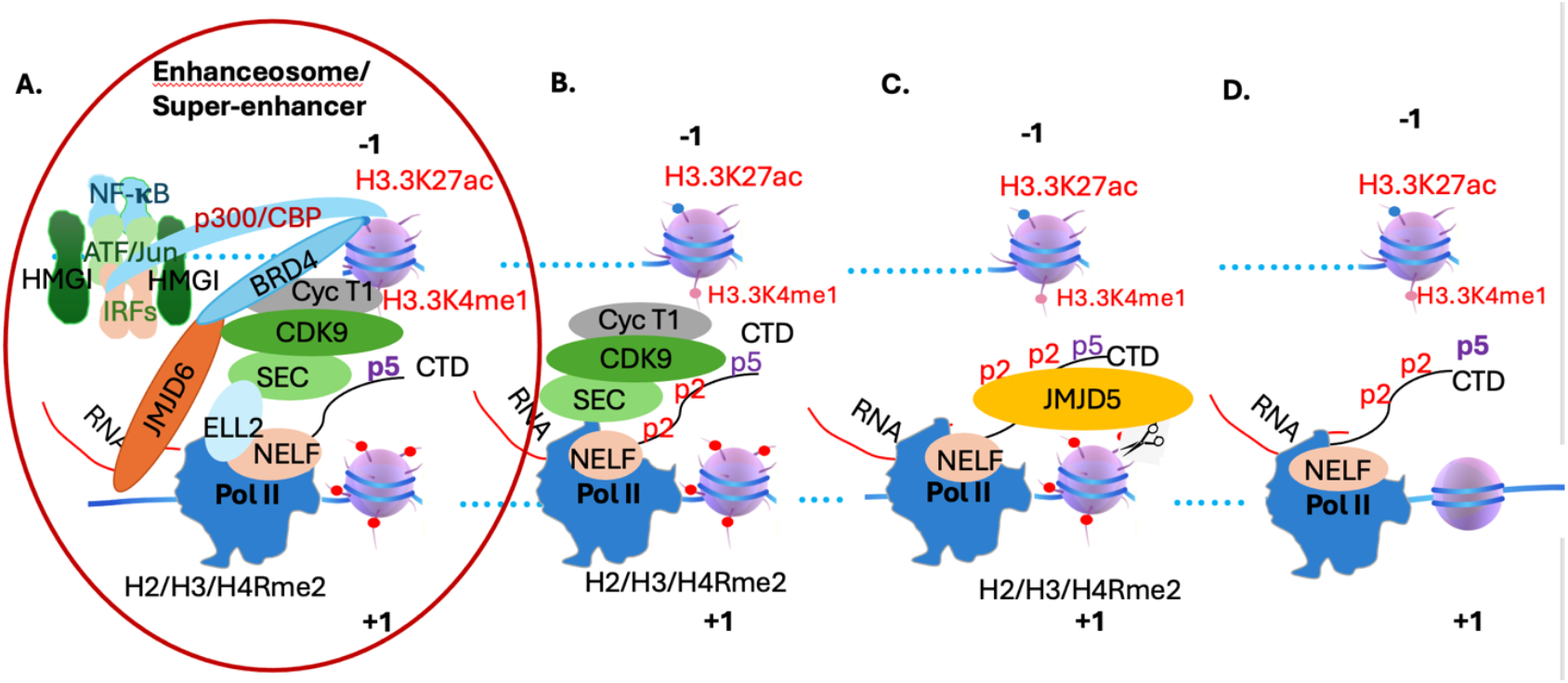
The coupling of the enhanceosomc and promoter-proximal Pol II pausing in metazoans. **A**. Virus infection triggers the activation of four major transcription factor families, which will recruit p300/CBP to the enhancer/promoter region of IFN-**β**. P300/CBP will generate a docking site to recruit BRD4, which in turn will recruit the CDK9 complex onto the super-elongation complex (SEC) with the help of JMJD6. The SEC will be recruited onto Pol II by ELL2. **B**. The CDK9 complex will generate a unique phosphorylation pattern on the CTD of Pol II, which will recruit JMJD5 via its CTD interaction domain. **C**. JMJD5 will cleave arginine-methylated histone tails of +1 nucleosome to generate a tailless nucleosome. **D**. Pol II will overcome the tailless +1 nucleosome and will be released from the pause status and engage in the elongation process.

## Materials and Methods

### Reagent

dBet6 and dCBP-1 were purchased from MedChemExpress with the Cat # HY-112588 and HY-134582, respectively. The chemical was dissolved in DMSO as a stock solution and stored at - 80°C. The rabbit antibody for Ser2ph-Ser2ph-Serp5-CTD of Rbp1 was a gift from Dr. David Bentley Lab as previously reported (Liu, Ramachandran et al. 2020). The Rpb1 antibody (sc-56767) and Actin antibody (sc-8432) were from Santa Cruz Biotechnology. The p300 antibody (#54062) is from Cell Signaling Technology.

### Western Blotting analysis

HEK 293 cell was seeded at 8*10^5 cells per well in the 6-well plate one night before treatment. dBet6 or dCBP-1 was added at the indicated concentration. After incubation, cells were harvested and lyzed in RIPA lysis buffer. The cell debris is discarded by centrifugation. Equal amount of extracted protein from the control and treatment groups is loaded on the SDS-PAGE gel for electrophoresis, and then the protein on the gel is transferred to the PVDF membrane and blotted with the indicated antibodies.

## Acknowledgments

We thank Drs. Philippa Marrack, Tony Gerber, and other members of National Jewish Health for their suggestions and support. Thank Dr. Richard Young for suggestions about the roles of condensates/phase separations in transcription regulation. Thank Dr. David Bentley’s group for the antibody specific for the unique phosphorylation pattern of CTD of Pol II. Thanks to Dr. Christopher Ott’s group for dCBP-1. The work is supported by an NIH grant GM135421 (G.Z.) and funds from NB Life Laboratory LLC, specifically private funds from Cheng-Yuan Zhang, Shi-Ning Xu, Lian-Hua Jin, Peng Sun, Yun-Xia Jiang, and Yongmei Jiang.

## Contributions

GZ conceived the concepts, designed experiments, conducted data analysis, and wrote up the manuscript. HL, LL, and QZ carried out the experiments. CO for dCBP-1 and suggestions.

